# Commonly-used FRET fluorophores promote collapse of an otherwise disordered protein

**DOI:** 10.1101/376632

**Authors:** Joshua A Riback, Micayla A Bowman, Adam M Zmyslowski, Kevin W Plaxco, Patricia L Clark, Tobin R Sosnick

## Abstract

The dimensions that unfolded proteins, including intrinsically disordered proteins (IDPs), adopt at low or no denaturant remains controversial. We recently developed an innovative analysis procedure for small-angle X-ray scattering (SAXS) profiles and found that even relatively hydrophobic IDPs remain nearly as expanded as the chemically denatured ensemble, rendering them significantly more expanded than generally inferred using fluorescence resonance energy transfer (FRET) measurements. We show here that fluorophores typical of those employed in FRET can contribute to this discrepancy. Specifically, we find that addition of Alexa488 to a normally expanded IDP causes contraction of its ensemble. In parallel, we also tested the recent suggestion that FRET and SAXS results can be reconciled if, in contrast to homopolymers, the radius of gyration (R_g_) of an unfolded protein chain can vary independently from its end-to-end distance (R_ee_). To do so, we developed an analysis procedure that can accurately extract both R_g_ and R_ee_ from SAXS profiles even if they are decoupled. Using this procedure, we find that R_g_ and R_ee_ remain tightly coupled even for heteropolymeric IDPs. We thus conclude that, when combined with improved analysis procedures for both SAXS and FRET, fluorophore-driven interactions are sufficient to explain the preponderance of existing data regarding the nature of polypeptide chains unfolded in the absence of denaturant.

---

Protein disorder is an essential component of diverse cellular processes (1-4). Unlike well-folded proteins, which populate a well-defined functional state, intrinsically disordered proteins (IDPs) and regions (IDRs) sample a broad ensemble of rapidly interconverting conformations (3-9) with biases that are poorly understood and difficult to measure. Of particular interest is the extent to which IDPs undergo compaction under physiological conditions (i.e., in the absence of denaturants). Such compaction would have broad implications for our understanding of protein folding, interactions and stability as well as the action of denaturants. Moreover, understanding the extent of collapse in disordered ensembles has profound implications for the development of realistic simulations of protein folding and interpretation of SAXS and FRET measurements (10, 11).

Our current understanding of the physiochemical principles that underlie whether a given polypeptide chain will fold, adopt a disordered but nevertheless compact ensemble, or behave as an expanded, fully-solvated self-avoiding random walk (SARW) is insufficient to explain existing data. Most of this understanding is derived from studies of proteins unfolded by high concentrations of chemical denaturants such as urea and guanidine hydrochloride (Gdn). Under these conditions the consensus is that proteins behave as SARWs, corresponding to a Flory exponent (*v*) of 0.60 in the relationship R_g_∝N^v^ (where R_g_ = radius of gyration and N = polypeptide chain length). In contrast, consensus is lacking regarding the behavior of disordered polypeptide chains at lower or no denaturant. Specifically, while numerous FRET (12-26) and computational studies (12, 15, 19, 24, 27-30) have argued that the expanded, disordered ensemble detected at high denaturant collapses upon transfer to low or no denaturant (ν < 0.5) (12, 15, 19, 24, 27-29, 31-36), almost an equal number of SAXS studies report no or only a minor contraction under these same conditions (11, 37-42).

A variety of recent studies have attempted to reconcile this crucial discrepancy (**Fig. 1A**). The application of more realistic simulations and analytical models, for example, results in FRET-derived distances having a smaller denaturant dependence (**Fig. 1A, bottom**) (41, 43-45). In parallel, improved SAXS data collection and analysis, including the use of the dimensionless Kratky plot to emphasize changes in v rather than R_g_, (which will increase upon addition of non-perturbative fluorophores near the ends of a chain due to the extra mass), likewise provides evidence for a minor contraction at guanidine hydrochloride (Gdn) concentrations below 2 M (**Fig. 1A, bottom**) (46, 47). Nevertheless, significant discrepancies persist in the absence of denaturant, even when the same approaches are used to analyze the same protein under identical conditions (**Figs. 1A-B**, **S1-S2**; **Movie S1**). Attempts have been made to reduce the appearance of the discrepancy by using a holistic analysis procedure (43, 44), but these analyses ignore its fundamental origins.

**Fig. 1.**
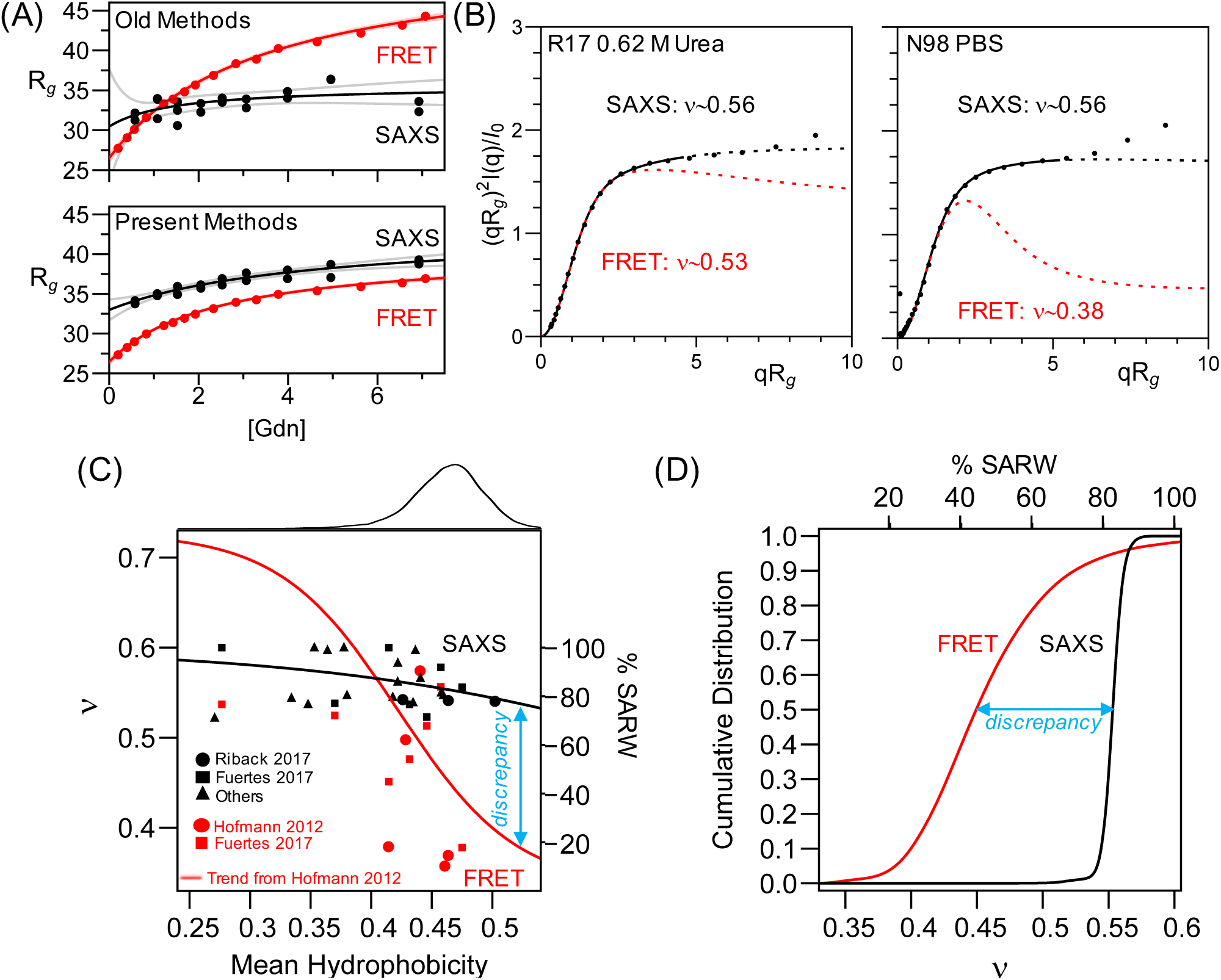
Improved analysis procedures do not eliminate discrepancy between SAXS and FRET-derived measurements of IDP dimensions. **(A)** R17 SAXS and FRET data (from (44)). Top, comparison of results obtained when FRET data is fit assuming a Gaussian chain and SAXS data is fit using the Guinier approximation. Bottom, SAXS and FRET data fit using our MFF analysis method and a similar approach (46). Black line is best fit hyperbolic trend line; grey lines are 95% confidence intervals. **(B)** SAXS profiles for R17 (left, from (44)) and N98 (right, from (43)) fit with the MFF are significantly different than the expected behavior using values of n taken from similar analysis of FRET data. Solid lines denote the region used in the fitting procedure; dashed lines represent extrapolation to higher values of q. Although ~500 points per scattering curve were fit, data shown were binned for presentation purposes only. The upturns or kinks in the data at higher values of qR_g_ are most likely due to errors in buffer subtraction, which is more challenging at high q, low sample concentration and/or reduced scattering contrast (e.g., at high denaturant, see Methods). **(C)** Trends of hydrophobicity (Kyte-Doolittle) versus n in the absence of denaturant derived from SAXS by applying the MFF to published data collected from foldable protein sequences (43, 46, 53-68). Also shown are results from FRET studies calculated as in (21) for published data (21, 43). Red trend line for FRET results is from (21). Black trend line is best fit to SAXS results shown. Top, histogram of hydrophobicity of representative proteins in the PDB (dataset from (46)). **(D)** Cumulative distributions of n for the representative proteins from the PDB, inferred from the trend lines shown in (C).

To comprehensively compare the results of SAXS and FRET, we collected published datasets for a variety of IDPs (**Fig. 1C-D**; **Table S3**). When analyzed using our simulations and molecular form factor (MFF), SAXS studies consistently find ν > 0.53 (mean = 0.55) whereas n derived from FRET studies typically falls below 0.50 (mean = 0.46). This 0.09 discrepancy is substantial relative to the entire range of n, which varies only from 0.6 (for a SARW) through 0.5 (where intra-chain interactions are equally favorable to solvent-chain interactions) to 0.33 (for a perfect sphere; it is somewhat higher for compact, non-spherical states). In general, SAXS results suggest that the conformational ensembles of a majority of unfolded proteins and IDPs with protein-like sequence composition are expanded, whereas FRET suggests otherwise (**Fig 1D**).

The above and other, similar results have led us and others to search for factors that might contribute to the persistent discrepancy between SAXS-and FRET-based views of IDP dimensions (10, 30, 40, 43, 44, 48-51). One alternative, herein denoted the “heteropolymer-decoupling hypothesis,” posits that the heteropolymeric nature of proteins leads to variation in the relationship between R_g_ and the polypeptide chain end-to-end distance (R_ee_), a relationship that is fixed at a ratio of 6.3 for a homopolymer adopting a SARW irrespective of chain length. Recent simulations, however, suggest that this ratio can vary significantly for heteropolymers (30, 40, 43, 44), with this “decoupling” offering a possible explanation for the discrepancy between SAXS (which is sensitive to R_g_) and FRET (which is sensitive to R_ee_). In contrast, a second hypothesis, herein denoted the “fluorophore-interaction hypothesis,” suggests that, in the absence of denaturant, the FRET fluorophores interact with each other and/or the polypeptide chain, causing the conformational ensemble of fluorophore-modified constructs to contract more than they would in the absence of these fluorophores (10, 46, 48, 51, 52).

Here we address the decoupling and fluorophore-interaction hypotheses. To achieve the latter we used SAXS to characterize the radius of gyration of IDPs in the presence and absence of fluorophore modifications. We find that labeling with fluorophores commonly used for FRET studies alters the conforma-tional ensemble populated in the absence of denaturant, decreasing its SAXS-measured dimensions by 10-20%. When coupled with improved analysis procedures employing realistic simulated ensembles for both SAXS and FRET, this fluorophore-induced collapse is sufficient to bring results from SAXS and FRET studies into agreement. In parallel, we present SAXS measurements on polyethylene glycol (PEG), confirming prior reports that the addition of fluorophores likewise causes the contraction of this otherwise SARW polymer (10), a finding that was recently questioned (43). Moreover, we show that SAXS can extract R_g_, v and R_ee_ with accuracies above 97% when analyzed using a new MFF developed for heteropolymers. These simulations are accurate enough to reproduce scattering data without the need to select only a sub-ensemble of conformations, as commonly used in other data fitting procedures. Finally, we demonstrate the extent that one can use small deviations from ideality in SAXS data to infer biases within the heteropolymer conformational ensemble.

## Results

### Fluorophore-labeling induces collapse of a highly expanded ensemble

To directly test the fluoro-phore-interaction hypothesis, we measured SAXS profiles of an unmodified IDP and the same IDP site-specifically modified with one or two copies of the commonly employed FRET fluorophore Alexa488. We chose this fluorophore because it is relatively small and hydrophilic, and thus considered unlikely to form interactions that would alter the conformational ensemble (44). As our test protein we used PNt, a well-behaved IDP comprising the amino terminal 334 residues of pertactin (69). To produce mono-and dual-fluorophore-modified PNt, we used a thiol-reactive Alexa488 to modify cysteine residues at either position 117 (PNtC-Alexa488) or positions 29 and 117 (PNtCC-Alexa488). As controls, we used the unmodified parent protein (PNt) and alkylation to produce constructs lacking fluorophores (PNtC-Alkd and PNtCC-Alkd).

The addition of Alexa488 reduces the SAXS-measured dimensions of PNt under both no-denaturant and intermediate denaturant conditions (**Fig. 2A, Table S1**). Specifically, R_g_ and v decrease nearly twice as much for the fluorophore-modified PNtCC-Alexa488 than for either PNtCC-Alkd or PNt (**Fig. 2B; Table S1**). These data indicate that the presence of Alexa488 leads to collapse of the PNt conformational ensemble. Of note, whereas 2 M Gdn is a good solvent (v > 0.50) for the unlabeled protein, fluorophore-labeling leads to measurable intramolecular interactions even at this relatively high denaturant concentration (**Fig. 2B, right**). The magnitude of this denaturant-dependent chain expansion is qualitatively similar to that observed by FRET for a variety of other proteins, consistent with a common origin (**Fig. 1B**) (43, 44). We also observed a fluorophore-dependent decrease in average R_g_ and v for the single-labeled construct PNtC-Alexa488 (**Fig. 2**), indicating that fluorophore-protein interactions are also significant. Of note, this collapse occurs despite steady-state fluorescence anisotropy values for PNtCC-Alexa488 of 0.11 and 0.08 in 0 and 2 M denaturant, respectively (**Table S2**), below the threshold typically considered as evidence of free rotation of the protein-attached fluorophores (43, 70). From this we conclude that addition of even a relatively small, hydrophilic fluorophore commonly employed for FRET measurements can significantly alter the dimensions of a disordered polypeptide chain (43, 44, 70).

**Fig. 2.**
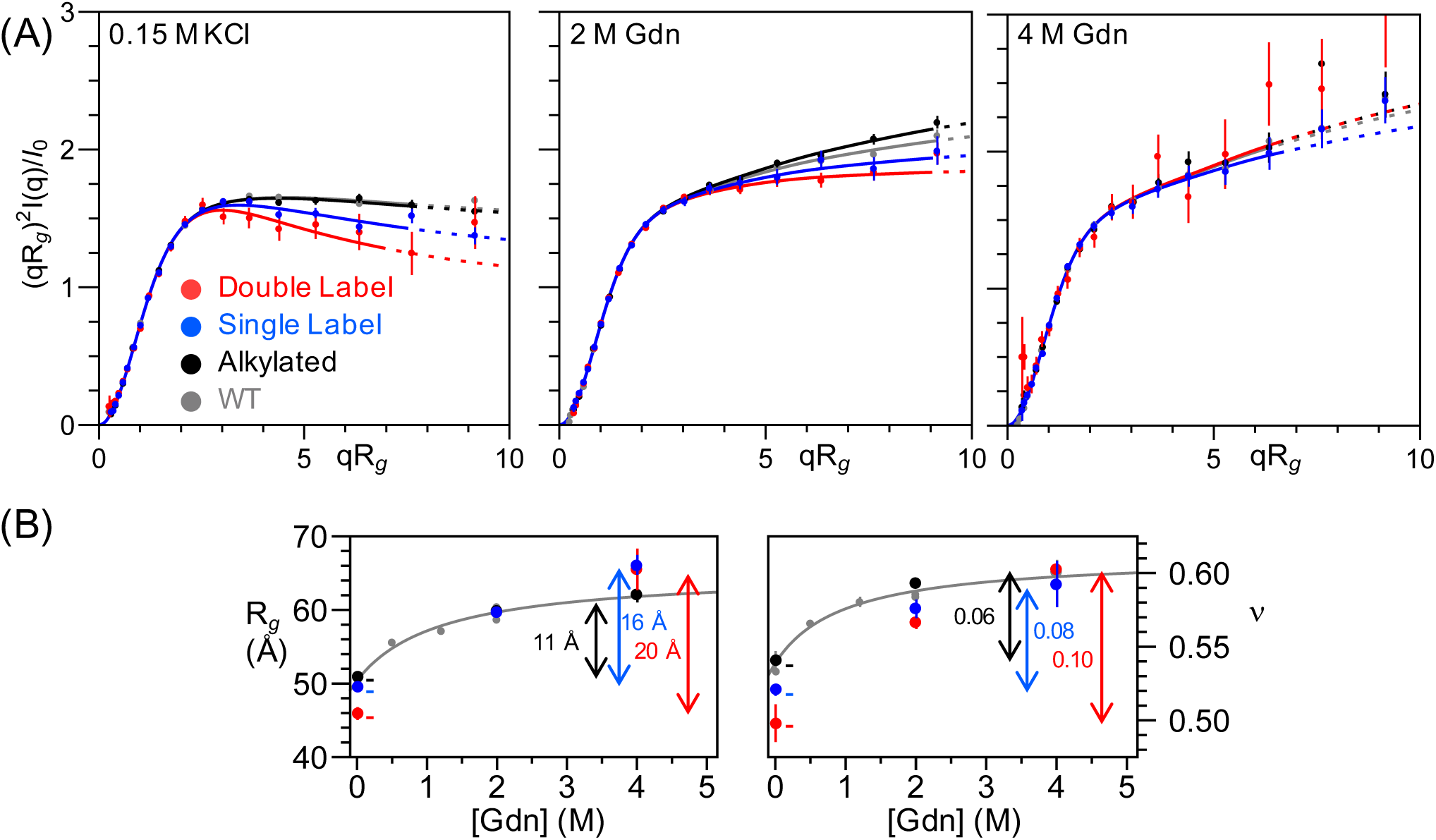
The addition of Alexa488 alters the scattering of PNt. (A) Dimensionless Kratky plots of wild type PNt (gray), PNtCC-Alkylated (black), PNtC-Alexa488 (single label, blue), and PNtCC-Alexa488 (double label, red) in 0.15 M KC1, 2 M Gdn, and 4 M Gdn. Error bars represent the propagated error from the standard deviation calculated assuming counting (Poisson) statistics where σ=√counts. Data were fit and displayed following the procedure described in Figure 1 (see also Methods). Results for alkylated PNt are indistinguishable from wild type PNt but there are significant differences for PNt labeled with Alexa488 fluorophores. (B) R_g_ and v as a function of Gdn concentration. Gray curves are reproduced from previous analysis of wild type PNt (46).

### The SAXS-derived dimensions of PEG are independent of polymer concentration

In an earlier study we reported that addition of Alexa488/594 to PEG resulted in a denaturant-dependent change in FRET (10), similar to that seen in FRET measurements of unfolded proteins. No contraction was observed, however, when the equivalent unlabeled polymer was studied using small angle neutron scattering. It has been proposed that the high (3 mM) concentrations of PEG used in this scattering study masked what would otherwise be a denaturant-dependent change in R_g_ (43). To test this, we measured SAXS profiles over a range of PEG and denaturant concentrations and found no evidence for a significant change in polymer dimensions (**Fig. 3**). Likewise, under all conditions we observe a Flory exponent of 0.60, further confirming that PEG behaves as a SARW independent of denaturant concentration. The fluorophore-interaction hypothesis thus remains the simplest interpretation of the denaturant-dependent changes in FRET observed for fluorophore-labeled PEG (10).

**Fig. 3.**
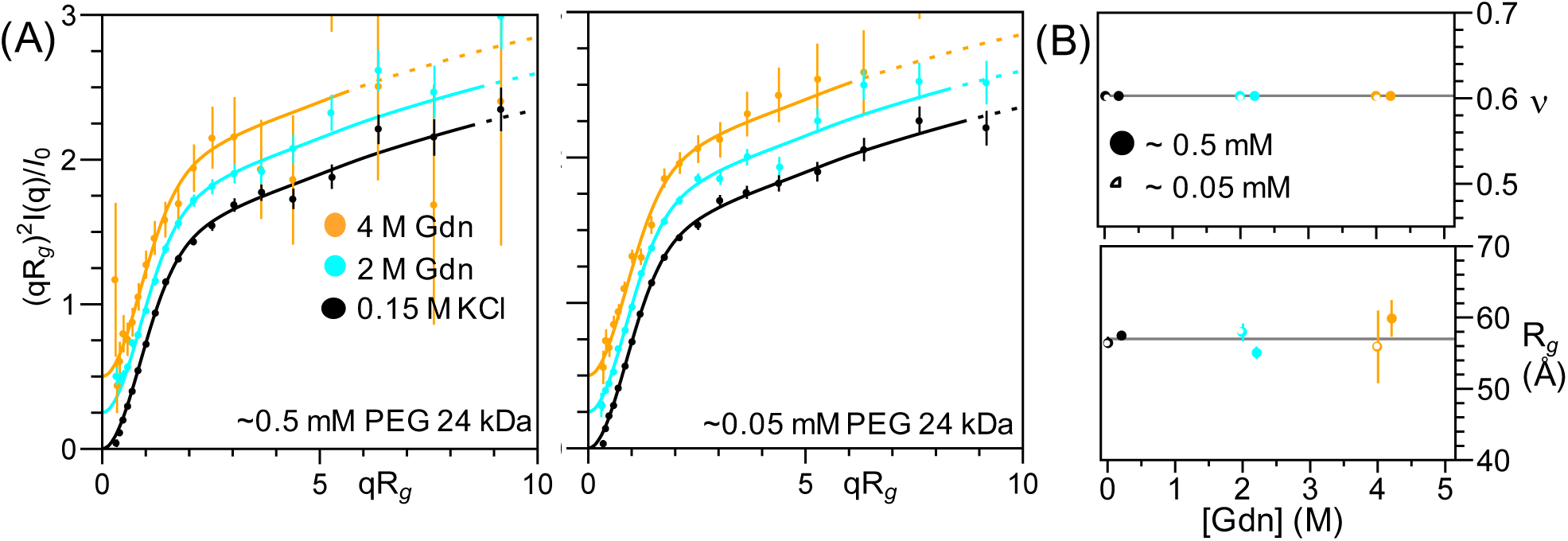
SAXS profiles of PEG are denaturant independent. **(A)** Dimensionless Kratky plots of 24 kDa PEG at 0.5 mM and 0.05 mM in 0.15 M KC1, 2 M Gdn, and 4 M Gdn. The normalized scattering profile of 24 kDa PEG is unchanged from 0-4 M Gdn over a broad range of PEG concentrations. Scattering profiles have been offset vertically for clarity. Data were fit and displayed using the procedure described in Figure 1. **(B)** R_g_ and v as functions of Gdn concentration for 0.5 mM and 0.05 mM PEG. Open and closed points are offset horizontally for clarity.

### Testing the heteropolymer-decoupling hypothesis

Taken together, the above observations indicate that fluorophores added to an IDP lead to significant contraction of its conformational ensemble, contributing to the different conclusions drawn from prior SAXS and FRET studies. These observations, however, do not rule out the possibility that heteropolymer-decoupling also contributes to the SAXS-FRET discrepancy.

To investigate whether and how decoupling between R_ee_ and R_g_ alter the SAXS profile and to test our ability to extract information from such deviations, we used *Upside*, our Cβ-level polypeptide chain simulation algorithm (71, 72), to simulate the scattering for unfolded ensembles of 50 protein sequences of 250-650 residues randomly chosen from the PDB. In its simplest form, *Upside* represents the polypeptide backbone with six atoms per residue (N, Cα, C, H, O, Cβ) and uses neighbor-dependent Ramachandran maps derived from a coil library (73). Such models are able to reproduce the R_g_ and NH residual dipolar couplings (RDCs) observed in unfolded proteins; these two parameters are sensitive to global and local properties of the backbone, respectively (74, 75). After creating an ensemble of backbone conformations, we add side chains (76) and determine scattering profiles of the hydrated versions (77). W e assigned each Cβ as either hydrophobic or polar (H/P), where the only favorable interactions are between Cβ atoms of aliphatic and/or aromatic residues. For each of the 50 sequences, 30 different β interaction strengths were used. These simulations yielded a range of deviations from G(v) = (R_ee_/R_g_)^2^ obtained from homopolymer simulations (Fig. S4A). From the 1500 resulting ensembles, we determined R_g_, v, and R_ee_ both directly from the atomic coordinates of the simulated ensemble and by fitting the hydrated SAXS profile (with added realistic random errors) of each ensemble using our MFF. The inferred end-to-end distance 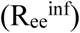 was determined using the relationship 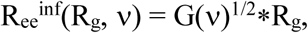, where G(v) was obtained from homopolymer simulations.

Compared to the true values calculated from the atomic coordinates, fits obtained using our MFF yield values of R_g_, v, and R_ee_ with a mean absolute deviation of only 1.3 Å, 0.011 and 4.2 Å, respectively, representing a 3%, 2% and 4% mean absolute error (**Fig S3**). The largest deviations are observed for more compact structures; for more extended conformations (v >0.54) the error is ~2%. To further reduce the small error associated with the application of our MFF derived from homopolymers to the scattering of heteropolymers we generated a new molecular form factor, MFF_het_, using the heteropolymer simulations described above. Application of this slightly modified MFF lowers errors in fitted R_g_, v, and R_ee_ to 0.5 Å, 0.005 and 2.7 Å, respectively, representing 1%, 1% and 2% mean absolute error (**Fig 4A-C**). These results demonstrate that SAXS analysis returns accurate values of R_ee_, in addition to R_g_ and v, indicating R_g_ and R_ee_ are coupled even for disordered heteropolymers.

**Fig. 4.**
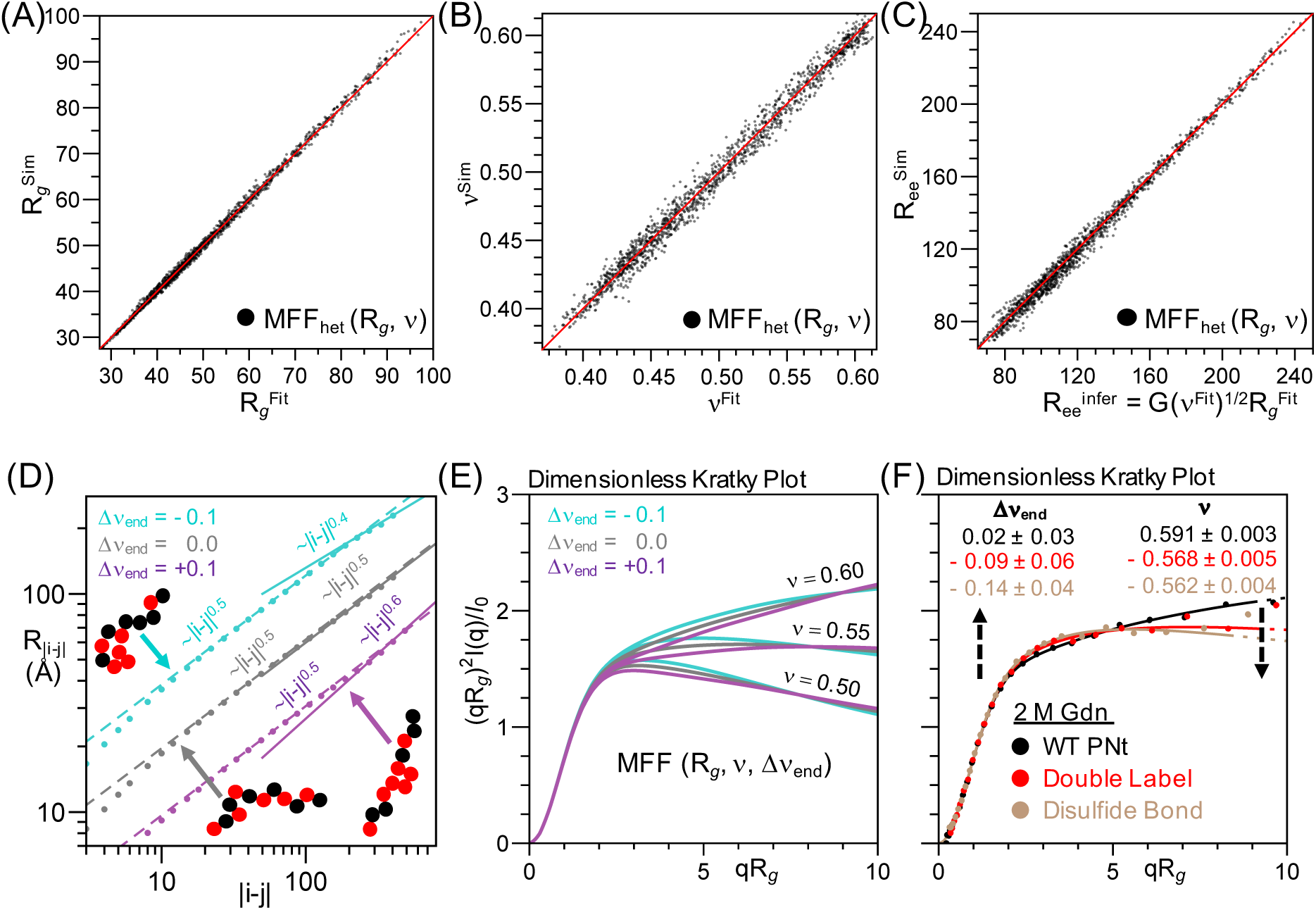
Fitting simulated SAXS data on realistic heteropolymer sequences demonstrates SAXS profiles are a robust measure of R_g_, v, and R_ee_ while also informing on the degree of heterogeneity. (**A-C**) Comparison of R_g_, v, and R_ee_ calculated from coordinates of HP-model simulations versus fit with our MFF_het_(R_g_, v) to SAXS profiles with randomly added experimental errors. **(D)** Deviations in v_ends_ are observed for heteropolymers with less well-mixed HP patterns (obtained by a fit to the slope of the dependence of the intra-chain distance, R_|i–j|_, on sequence separation, |i–j| where |i–j| > N/2). **(E)** Effects of Δv_end_ at different values of v. **(F)** Experimental data fit to MFF(R_g_,v, Δv_end_) demonstrates fluor-ophore labeling and loop formation via disulfide bonds in PNt induces significant and measurable deviations.

We next considered whether our conclusions are sensitive to the details of our model or energy function. To test this, we conducted additional simulations using a more detailed version of the *Upside* algorithm that is capable of *de novo* folding of proteins shorter than 100 residues (71, 72). In this version, each of the 20 side chains is represented by a multi-position eccentric bead that allows for detailed packing of the core. The energy function includes hydrogen bonds, side chain-side chain and side chain-backbone interactions, amino acid-dependent dihedral angle potentials and a desolvation term. Using this model, we generated 30 ensembles for each of six proteins (PNt and five other proteins randomly selected from the list of 50 above), using short simulations that sample only the unfolded state. Ensembles were obtained by running replica exchange simulations over a temperature range from 280-320 K, as described previously (71, 72). Values of v obtained from these ensembles ranged from 0.4 to 0.6, depending on the simulation temperature. Significantly, values of R_ee_, R_g_ and v obtained from these ensembles are in close agreement with values determined using our MFF, with nearly the same accuracy (**Fig. S3**). Hence our conclusion that R_g_ and R_ee_ remain coupled even for heteropolymers is thus robust to the details of the simulations.

### Measuring deviations from ideality in heteropolymers

MFF_het_ accurately captures the overall dimensions of disordered heteropolymers. Nevertheless, small but measurable deviations are observed for proteins in our test set with less well-mixed HP patterns (**Fig. S4**). These differences can be seen in the in-tra-molecular distance distribution plot, where the slope at separation distances |i-j|>N/2 can be different than the average slope, which defines the global v value (**Fig. 4D**). We define change in slope as Δv_end_ (**Fig. 4D**). Negative values of Δv_end_ correlate with a preponderance of hydrophobic residues at the ends of the polypeptide sequence (**Figs. 4D, S4C**) and with deviations in G(v) (**Fig. S4A**) (R^2^~0.84). The SAXS profile is most sensitive to ΔVend at low qR_g_ (**Fig. 4E**).

To extract this extra information from the SAXS data, we generated a more general, three-parameter form factor, MFF(R_g_, v, Δv_end_) (**Fig. 4E-F**; **Movie S2**). To demonstrate its ability to yield useful infor-mation, we fit data from PNt, PNtCC-Alexa488 and a circularized (disulfide-bonded) PNtCC at 2 M Gdn (**Fig. 4F**). Δv_end_ decreases from ~ 0 for PNt to ~ −0.1 for PNtCC-Alexa488 and PNtCC, consistent with the increase in interactions at the amino-terminus of the chain. These data demonstrate that for disordered polymers, SAXS is sensitive to sequence-dependent deviations from homopolymer behavior (**Fig. 4E-F**) while still able to accurately measure R_g_ and v (**Fig. 4A-C**).

## Discussion

Whereas SAXS measurements point to water being a good solvent (v > 0.54) for unfolded polypeptides, FRET measurements typically find the opposite (v < 0.50). We find here, however, that improved analysis procedures and fluorophore-fluorophore and/or fluorophore-chain interactions are sufficient to explain this discrepancy. Thus, the fundamental and significant conclusion remains that, even in the absence of denaturant, water is a good solvent for most polypeptide sequences. Specifically, we find that labeling with Alexa488, a fluorophore commonly used for FRET measurements, can alter the conforma-tional ensemble of a disordered protein, decreasing R_g_ and v even when the fluorescence anisotropy is low relative to accepted limits for free fluorophore rotation (43, 70). In combination with prior studies (10), similar conclusions can be inferred for PEG, a known SARW. These findings, along with our prior result that disordered chains undergo a mild expansion in denaturant (46) and improved methods for extracting R_g_ values from FRET data (41, 43-45), now provide a sufficient framework for resolving discrepancies between SAXS and FRET on the dimensions of disordered proteins.

Consistent with our findings of fluorophore-induced effects, others have found that molecular dimensions inferred by FRET can be dependent on the fluorophore pair used, with more hydrophobic fluorophores leading to smaller (more compact) dimensions (44). All-atom molecular dynamics simulations with a Alexa488/594 fluorophore pair, for example, resulted in a 10% contraction of an IDP even in 1 M urea (78). Likewise, a recent study found that smFRET signals from both DNA and PEG, which are often referred to as “spectroscopic rulers,” are dependent on solvent conditions under which the dimensions of the chains were expected to be invariant (52).

However, in apparent disagreement with our data, Fuertes *et al.* (43) conducted SAXS measurements on five IDPs with and without Alexa488/594 and concluded that, *on average*, the alterations seen upon the addition of fluorophores were minimal. When considered for each protein separately, however, the differences appear significant relative to the narrow range of possible values. Specifically, for the five proteins characterized in that study, v_unlabel_-v_label_ =0.08, 0.03, 0.03, −0.02, −0.04 (or 0.09, 0.06, 0.03, −0.02, −0.08 when analyzed using our procedures; Fig. S5). Although Fuertes *et al.* assert that only one protein (NLS), exhibits fluorophore-induced contraction (50), in fact four of the five proteins they tested had statistically significant fluorophore-induced changes in n, with more than half exhibiting a fluorophore-induced contraction (43) of similar magnitude to the contraction we observed for fluorophore-labeled PNt in water (46) (**Fig. S5**). Together, these data support a consistent picture of fluorophore-induced perturbations, contributing to differences in the magnitude and denaturant dependence of R_g_ inferred from SAXS and FRET.

The other factor that has been suggested to contribute to the discrepancy between SAXS and FRET results is deviations from the proportional relationship between R_g_ and R_ee_ that arise when analyzing heteropolymers versus homopolymers (43). Underlying this view is the observation that, if one reweights the ensemble (i.e., calculates R_g_ using only a subset of conformations), many possible values of R_ee_ are consistent with any given R_g_ (and vice-versa). Rather than selecting a sub-ensemble of conformations to fit a couple of parameters, we have taken an alternative approach (46). We generate physically plausible ensembles at the outset, create a MFF using these entire ensembles, and examine whether it fits the data in its entirety. We find that our MFF accurately matches the entire scattering profile (rather than just the R_g_), which provides strong support for our procedure. Since we can calculated the values of R_g_ and R_ee_ directly from the underlying ensembles, we have a procedure to obtain these two parameters by fitting the SAXS data with our MFF.

This MFF is imperfect in the sense that slightly different ensembles can be fit using the same R_g_ and v parameters. But the error is very low for these two parameters relative to their true values (**Fig. 4A-C**). Inclusion of heteropolymer effects does not alter this conclusion. From these results, we conclude that SAXS is well suited to extract both R_g_ and R_ee_ for disordered heteropolymers, while circumventing potential artifacts due to fluorophore interactions with polypeptide chains. This conclusion does not negate the potential of FRET to measure dynamics, binding and conformational changes; it does, however, emphasize that caution must be exercised when employing FRET to infer quantitative distances in the original, unlabeled biomolecule.

Nearly a dozen IDP SAXS datasets reported here and previously (46) have been shown to fit well to our general MFF (**Tables S1, S3**). This finding suggests that the interactions that drive chain contraction are spread along protein sequences. Water-soluble, well-folded protein sequences tend to be well-mixed heteropolymers, with relatively small stretches of consecutive hydrophobic residues (79). These well-mixed sequences tend to behave as homopolymers when measured by global, low resolution methods such as SAXS. Indeed, we have demonstrated that, with sufficient data quality, poorly mixed sequences can be identified by their deviation from our MFF (**Fig. 4D-F**). Larger deviations can occur for some IDPs, especially those with partial folding, unusual sequence patterning (e.g., block copolymers) and/or under crowded conditions that may serve specific functions (80, 81).

Our results reinforce the view that water is a good solvent for most unfolded polypeptides, a property that should reduce misfolding and aggregation while simultaneously facilitating synthesis and transport. That most proteins nevertheless readily fold in water suggests that the interactions that drive folding are more stabilizing - i.e., more easily overcome the ability of water to solvate the unfolded state - than those that promote non-specific collapse. Indeed, the observation that, despite the minimal evidence of significant unfolded-state contraction even in the complete absence of denaturant, some proteins remain stably folded in up to 6 M Gdn (42, 82) suggests that native interactions are far more favorable than any non-specific interactions associated with collapse. Given, however, the highly specific nature of the interactions formed in native proteins, their preferential ability to overcome the solvation of the unfolded chain is perhaps not surprising.

## Materials and Methods

### Protein purification

PNtCC and PNtC were expressed in *E. coli* BL21(DE3)pLysS and purified from inclusion bodies as described previously (46, 69, 83), with the following modifications. After inclusion body solubilization, PNt constructs were refolded in 50 mM Tris pH 7.2 with 50 mM ß-mercaptoethanol (ßME). Prior to the final size exclusion chromatography step, 20 mM ßME was added to the protein stock solution.

### Alkylation

Purified PNtCC or PNtC (70 uM) in 50 mM Tris pH 8, 5 mM EDTA was reduced with 5 mM TCEP for 30 min at room temperature while stirring. Alkylation was initiated by addition of 10 mM iodoacetamide in water. PNt constructs were incubated for 30 min at room temperature while stirring in the dark. The alkylation reaction was quenched with the addition of 20 mM fresh DTT. Excess reagents were removed by size exclusion chromatography (Superdex 16/60 S200). Alkylation efficiency (100%) was determined by mass spectrometry (MS).

### Alexa488 labeling

PNt constructs were reduced as above. Alexa488 C5 maleimide (ThermoFisher Scientific) was resuspended in DMSO to 5 mg/ml and added to the reduced protein dropwise while stirring, to a final ratio of 5:1 fluorophore:cysteine. The reaction proceeded overnight at 4°C while stirring in the dark. Free fluorophore was separated from fluorophore-labeled protein by size-exclusion chromatography (Superdex 16/60 S200), protected from light at all times. Labeling efficiency (100% for PNtCC; >50% for PNtC) was determined by mass spectrometry.

### Steady-state anisotropy

Steady-state anisotropy measurements were performed on a QM-6 T-format fluorometer (Horiba) at room temperature in 50 mM Tris pH 7.5 and 2 M Gdn where indicated. Samples containing 1 uM Alexa-488 were excited with 494 nm vertically polarized light. Anisotropy (r) was calculated using the emission at 516 nm by the equation

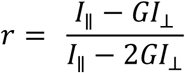

Where G is the instrument correction factor. G was calculated from the emission of free-Alexa488 at 516 nm after excitation with 494 nm horizontally polarized light by the equation

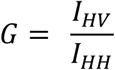

### SAXS

Data were collected at the BioCAT beamline at the Advanced Photon Source (Argonne National Lab) using a GE Lifesciences Superdex 200 SEC column with the scattering presented as q=4π sinθ/γ, where 2θ is the scattering angle and γ is the X-ray wavelength (1 Å). Error bars for experimental data points are based on counting (Poisson) statistics, where the standard deviation is given by σ=√(photons) because the X-ray detector counts individual photons (84). During the averaging and buffer subtraction steps, error is propagated in quadrature (e.g., for X=aU+bV, σX^2^ = (aσU)^2^ + (bσV)^2^ where a and b are constants). When I(q) is plotted, the error bar is the standard deviation value scaled for instrumental factors such as X-ray flux, solid angle of the detector, etc. as appropriate for propagation of errors (for x=au, σ_x_=aσ_U_, when a is a constant). Similarly, when data are presented in the dimensionless Kratky plot, the value of σ is scaled by the value of qR_g_^2^/I_o_ obtained from fitting the data. Approximately 1,000 data points are collected for each scattering curve. For presentation purposes only, after the fitting procedure (below), data points are binned to maintain clarity and facilitate comparisons between scattering curves. The standard deviation for the binned data was obtained by adding errors in quadrature (84).

### Data fitting

Buffer subtraction is more challenging at high q (due to hydration effects) and at high denaturant concentrations (due to reduced scattering contrast). For these reasons, and to maintain consistent criteria with our earlier study (46), we fit all data only to q=0.15 or 0.10 Å^−1^, in water or elevated denaturant (above 3 M), respectively. Solid lines denote the region used in the fit while dashed lines represent an extrapolation to higher values of q. R_g_ and v were obtained by fitting the data I(q) versus q using a standard non-linear least squares procedure (*χ*^2^ minimization with data weighted by 1/σ^2^ in Mathematica) to the indicated MFF. This procedure is appropriate as the error is the expected measurement standard deviation (84) obtained based on counting statistics with σ^2^=counts. This fitting procedure resulted in a reduced *χ*^2^ near 1.0, as expected for a high quality fit with proper error analysis.

### Simulations and generation of MFF

Calculations were performed at the University of Chicago Research Computing Center using a version of our *Upside* molecular dynamics program (71, 72) modified for Cβ-level interactions, as done previously (46). After creating an ensemble of backbone conformations, we add full side chains with the TreePack algorithm (76) and determine scattering profiles of the hydrated versions using the FOXS program (77). To calculate FRET efficiencies we used our MFF and previously published simulations (46) to determine a dimensionless FRET function, E(<R_ee_^2^>^0.5^/ R_0_,v), where R_0_ is the Förster radius, and used the expected dependence between R_g_, R_ee_, v, and N. Comparison between our SAXS/FRET methods and others are shown in Figs. S1-2.

### Code for simulation and analysis

Code and associated files necessary to produce simulations and all SAXS and FRET data analysis herein can be accessed at http://www.(to be added prior to publication). Additionally, our webserver http://sosnick.uchicago.edu/SAXSonIDPs is available for fitting SAXS data with our MFF(v,R_g_).

## Acknowledgements

We thank Srinivas Chakravarthy for assistance in the SAXS measurements and Matthew Champion for assistance with the mass spectrometry experiments. This work was supported by National Institutes of Health (NIH) Research Grant R01 GM055694 (TRS), the W. M. Keck Foundation (PLC) and NSF grants GRF DGE-1144082 (JAR) and MCB 1516959 (CR Matthews). Use of the Advanced Photon Source, an Office of Science User Facility, operated for the Department of Energy (DOE) Office of Science by Argonne National Laboratory, was supported by the DOE under Contract No. DEAC02-06CH11357. This project was supported by the NIH (2P41RR008630-18 and 9 P41 GM103622-18).

## Supplementary Information for

**Includes:**

Supplementary Figures (S1-S5)

Supplementary Tables (S1-S3)

Supplementary Movies (S1-S2)

**Figure S1:**
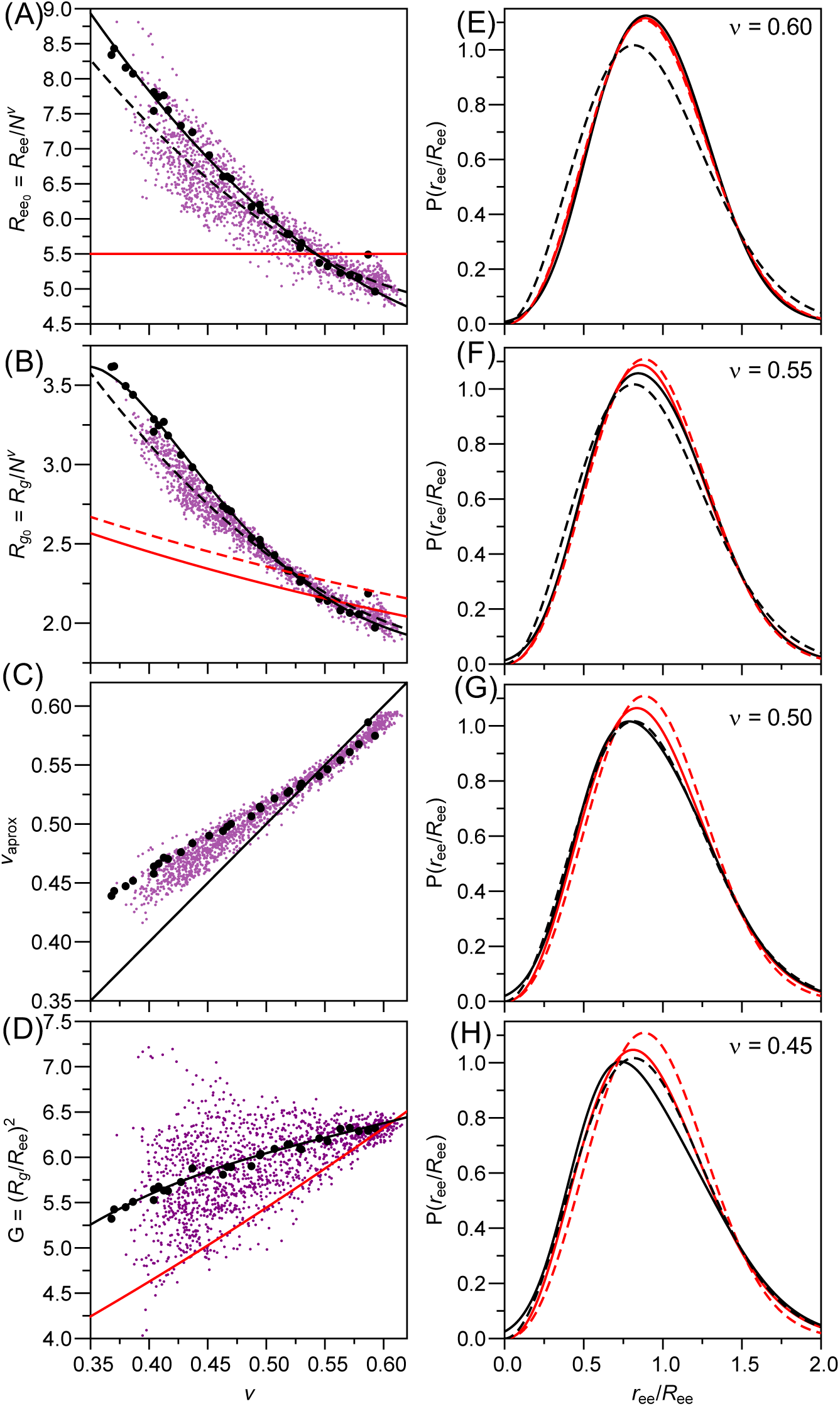
Comparison of polymer models used in SAXS and FRET. (**A-D**) Comparison of ν vs. various polymer quantities. Black dots and solid lines, ALL-H PNt simulations and resulting trend from Riback *et al.* 2017 (46). Purple dots, values from FIP-model simulations. Dashed black lines, trends from FIP-model simulations with near-even FIP distributions. Red line, trends from Borgia *et al.* 2016 (44). **(B)** Dashed red line, trend from Hofmann *et al.* 2012 (21). **(C)** Comparison of ν determined from the R_|i.j|_ plot and v approximated assuming R_0_=5.5 and the procedure in (21). Black line is an x=y line for reference. **(D)** Red line, trend from Zheng *et al.* 2018 (45). (**E-H**) Dimensionless (r_ee_/R_ee_) end-to-end distribution. Red dashed, SARW model; black dashed, Gaussian model; solid red, model from Zheng *et al.* 2018 (45). Solid black, model presented herein based on ALF-H PNt simulations at the noted v values.

**Figure S2:**
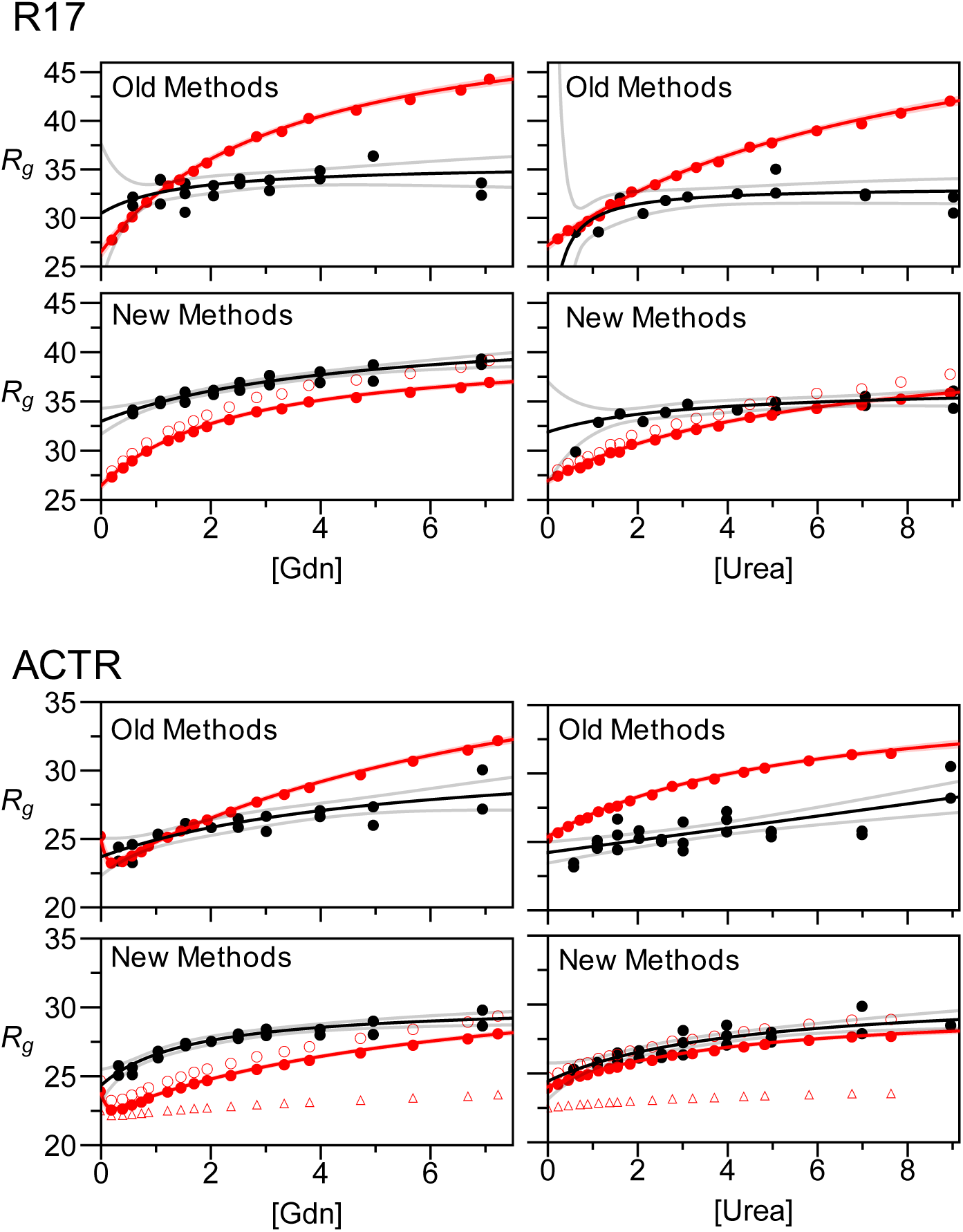
Comparison of published SAXS and FRET data on the same systems. Legend equivalent to Figure 1A. R17 and ACTR SAXS and FRET data (from Borgia *et al.* 2016 (44)). Top, ‘Old Methods’ Comparison between FRET data fit assuming a Gaussian chain and SAXS data fit with the Guinier approximation. Bottom (filled circles), ‘New Methods’ SAXS and FRET fit with MFF from (46). Lines shown are best fit hyperbolic trend and mean prediction lines in opaque and slightly transparent, respectively. Open circles are models from Zheng *et al.* 2018 (45). Open triangles correspond to Song *et al.* 2017 (40) determined at http://dice.utm.utoronto.ca.

**Figure S3:**
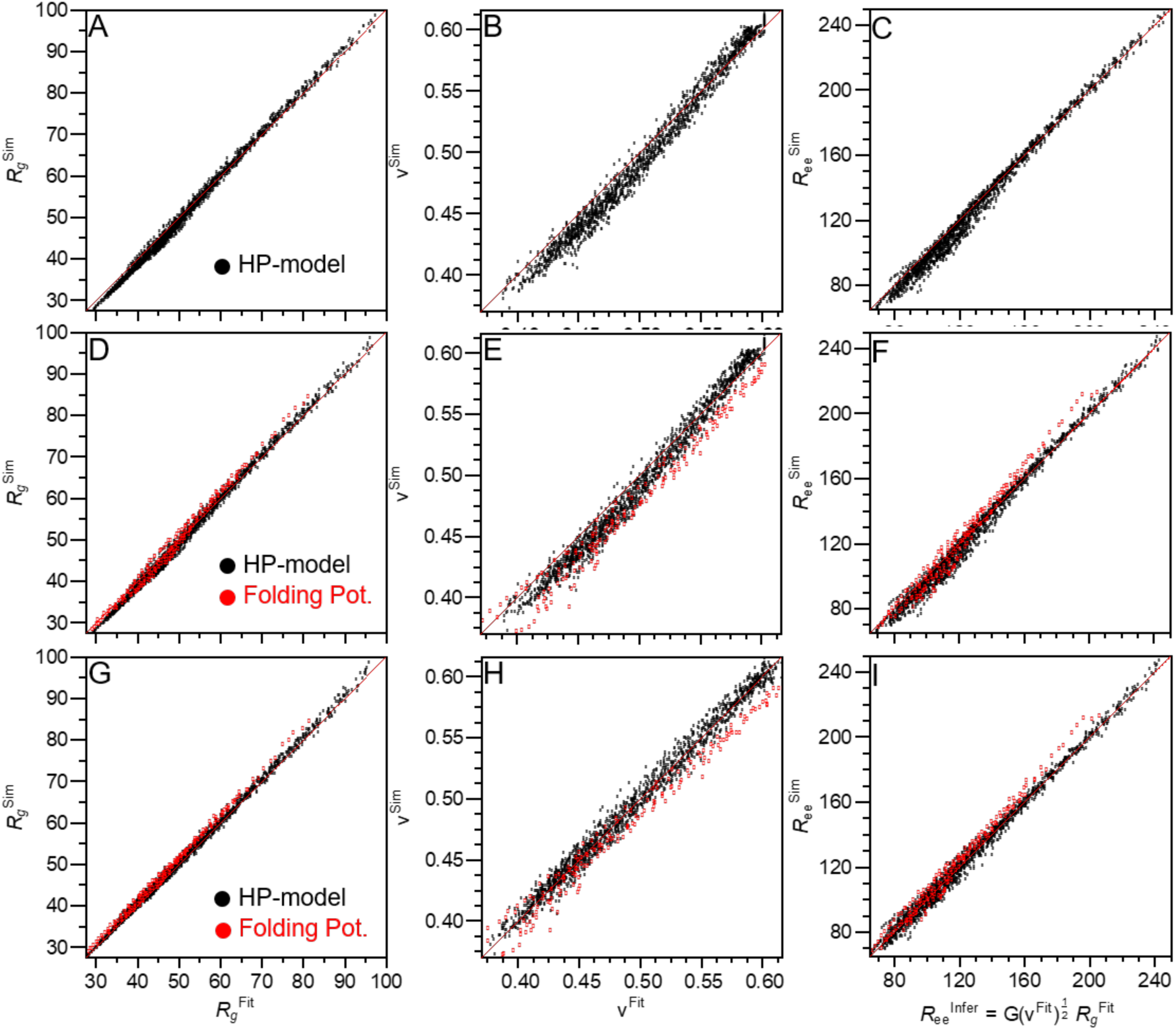
Comparison between parameters obtained from the two-parameter MFFs and true values for the conformational ensembles generated using *Upside* simulations (72). (**A-C**) Comparison of R_g_, v, and R_ee_ calculated from coordinates of HP-model simulations versus fit with our published MFF(R_g_, v) to SAXS profiles with randomly added experimental errors. (**D-I**) Same as panels (A-C) but for simulations using the advanced *Upside* energy function and fit with (**D-F**) our original MFF(R_g_, v) or (**G-I**) the new MFF_het_(R_g_, v).

**Figure S4:**
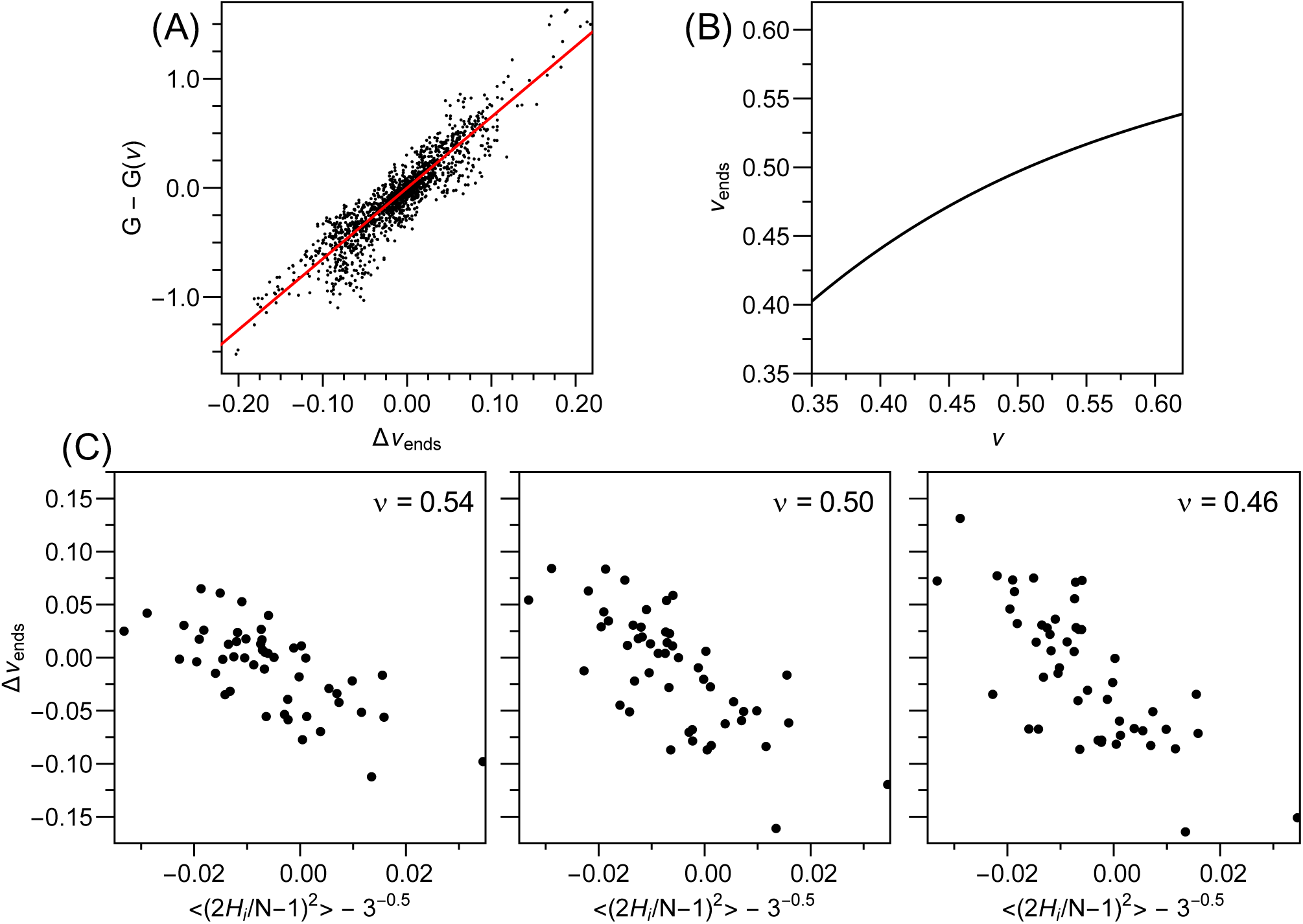
Heteropolymer simulations. (**A**) Correlation between Δv_end_ with G-G(v) in HP-model simulations ((G=R_ee_/R_g_)^2^) where G(v) is the G value expected for a homopolymer at that value of v. Red line is the best linear fit. (**B**) Dependence on v_ends_ with v extrapolated to simulations where G=G(v). (**C**) Dependence on Δv_end_ with the average mean squared deviation of hydrophobic residues from the center of the chain. The x-axis is referenced such that an evenly spaced heteropolymer has a zero x value. The v values for the three conditions are shown.

**Figure S5:**
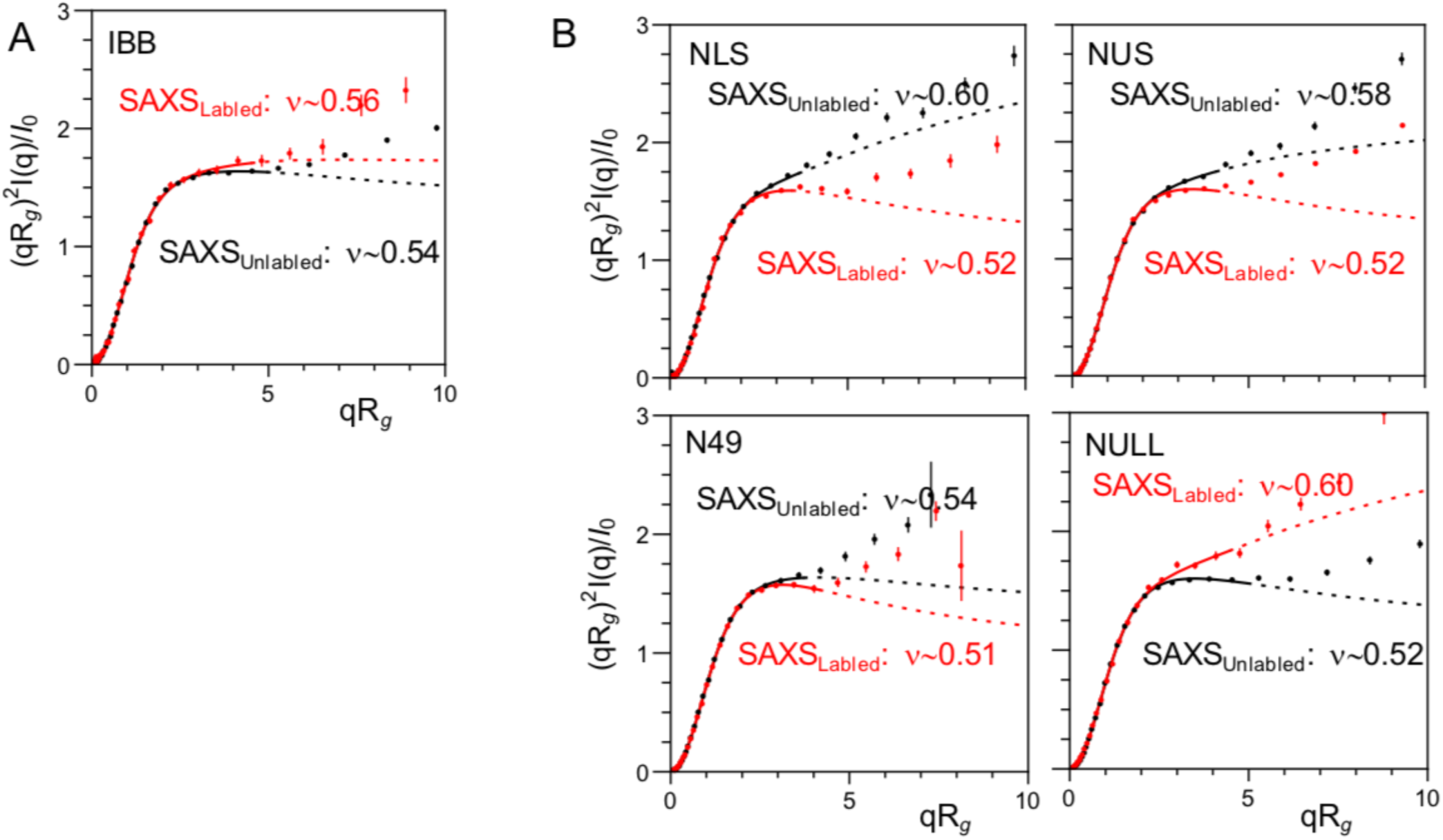
Effects of FRET fluorophores on the conformational ensemble of other IDPs. Scattering data from Fuertes *et al.* (43) of five IDPs with and without Alexa488/594 labels fit using the MFF from (46). (**A**) Data with labeled and unlabeled v values within one standard deviation fit error (indistinguishable). (**B**) Data with labeled and unlabeled proteins v values with greater than one standard deviation fit error (distinguishable).

**Table S1:** SAXS on PNt; data fit to MFF_homopolymer_.

|  |  | 0.150 M KCl | 2 M Gdn | 4 M Gdn |
| --- | --- | --- | --- | --- |
| PNt | $R_g$ (Å): | $51.1 \pm 0.1$ | $58.7 \pm 0.1$ | $62.2 \pm 0.5$ |
| | $v$: | $0.542 \pm 0.002$ | $0.584 \pm 0.001$ | $0.601 \pm 0.008$ |
| | $\chi^2_r$: | 1.11 | 0.986 | 0.946 |
| PNtCC-Alkd | $R_g$ (Å): | $51.0 \pm 0.2$ | $59.8 \pm 0.2$ | $62.2 \pm 1.0$ |
| | $v$: | $0.541 \pm 0.003$ | $0.593 \pm 0.002$ | $0.603 \pm \text{N/A}$ |
| | $\chi^2_r$: | 1.00 | 1.04 | 1.10 |
| PNtC-Alkd | $R_g$ (Å): | $51.8 \pm 0.2$ | $60.1 \pm 0.5$ | $62.0 \pm 0.8$ |
| | $v$: | $0.527 \pm 0.002$ | $0.588 \pm 0.004$ | $0.603 \pm \text{N/A}$ |
| | $\chi^2_r$: | 1.11 | 0.977 | 0.996 |
| PNtCC-Alexa488 | $R_g$ (Å): | $45.9 \pm 0.7$ | $59.6 \pm 0.4$ | $65.5 \pm 2.7$ |
| | $v$: | $0.498 \pm 0.012$ | $0.566 \pm 0.004$ | $0.603 \pm \text{N/A}$ |
| | $\chi^2_r$: | 0.995 | 0.975 | 0.867 |
| PNtC-Alexa488 | $R_g$ (Å): | $49.6 \pm 0.2$ | $59.8 \pm 0.6$ | $66.1 \pm 1.2$ |
| | $v$: | $0.521 \pm 0.003$ | $0.576 \pm 0.006$ | $0.593 \pm 0.015$ |
| | $\chi^2_r$: | 0.892 | 1.03 | 1.02 |

**Table S2:** Alexa488 fluorescence anisotropy and lifetime measurements^1^.

|  | [Gdn] (M) | Anisotropy | Lifetime (ns) |
| --- | --- | --- | --- |
| Free Alexa488 | 0 | $0.017 \pm 0.001$ | 4.72 |
| | 2 | $0.0304 \pm 0.0007$ | 4.82 |
| PNtC-Alexa488 | 0 | 0.132 (n=2) | 4.67 |
|  | 2 | 0.071 (n=2) | 4.58 |
| PNtCC-Alexa488 | 0 | $0.109 \pm 0.005$ | 4.56 |
| | 2 | $0.074 \pm 0.001$ | 4.59 |

<sup>1</sup>Error shown is the SEM of 3 measurements.

**Table S3:** Analysis of SAXS data on published IDPs with our MFF^1^.

| Protein | Citation | $R_g(\text{\AA})$ | error | n | errors | $c_r^2$ | q min | q max |
| --- | --- | --- | --- | --- | --- | --- | --- | --- |
| ID3 of CBP | (54) | 75.7 | 0.3 | 0.586 | 0.001 | 2.44 | 0.01 | 0.15 |
| Ash1 | (55) | 29.2 | 0.2 | 0.603 | 0.007 | 1.04 | 0.01 | 0.15 |
| Met2 | (56) | 52.7 | 0.1 | 0.542 | 0.002 | 0.98 | 0.01 | 0.15 |
| NSP | (43) | 37.3 | 0.2 | 0.603 | 0.006 | 1.15 | 0.01 | 0.1 |
| N98 | (43) | 34.3 | 0.1 | 0.556 | 0.002 | 2.62 | 0.03 | 0.15 |
| IBB | (43) | 33.2 | 0.1 | 0.538 | 0.002 | 1.05 | 0.01 | 0.15 |
| NUS | (43) | 27.3 | 0.1 | 0.578 | 0.010 | 1.07 | 0.01 | 0.15 |
| NLS | (43) | 24.3 | 0.1 | 0.603 | 0.006 | 0.59 | 0.01 | 0.15 |
| N49 | (43) | 16.6 | 0.1 | 0.537 | 0.012 | 0.98 | 0.01 | 0.225 |
| NUL | (43) | 33.3 | 0.1 | 0.523 | 0.004 | 2.20 | 0.01 | 0.15 |
| Msh6 NTR | (57) | 61.4 | 0.2 | 0.547 | 0.001 | 1.93 | 0.01 | 0.15 |
| HMPV (P1-60) | (58) | 26.5 | 0.1 | 0.603 | 0.003 | 0.80 | 0.01 | 0.15 |
| redAFP | (53) | 22.2 | 0.1 | 0.550 | 0.006 | 1.00 | 0.01 | 0.225 |
| MeCP2 | (59, 60) | 59.7 | 0.9 | 0.550 | 0.002 | 1.11 | 0.02 | 0.15 |
| Ki-1/57 | (59, 62) | 49.5 | 0.7 | 0.539 | 0.003 | 0.34 | 0.01 | 0.15 |
| CSD1 | (59, 61) | 32.9 | 0.5 | 0.565 | 0.011 | 0.50 | 0.02 | 0.15 |
| HrpO | (59, 63) | 35.3 | 0.3 | 0.603 | N/A | 0.72 | 0.01 | 0.15 |
| II-1 | (59, 65) | 38.7 | 0.6 | 0.525 | 0.005 | 0.42 | 0.013 | 0.142 |
| ERM (1-122) | (59, 64) | 37.5 | 0.5 | 0.548 | 0.002 | 0.14 | 0.013 | 0.147 |
| FEZ1 | (59, 66) | 35.2 | 0.1 | 0.603 | N/A | 0.16 | 0.01 | 0.15 |
| p53 | (59, 67) | 29.1 | 0.1 | 0.569 | 0.007 | 0.29 | 0.01 | 0.15 |
| PIR | (59, 68) | 26.5 | 0.2 | 0.553 | 0.008 | 0.10 | 0.01 | 0.15 |

<sup>1</sup>q is in units of inverse angstroms. When (59) is included in citation column, it provided data obtained from other studies.

**Movie S1:**
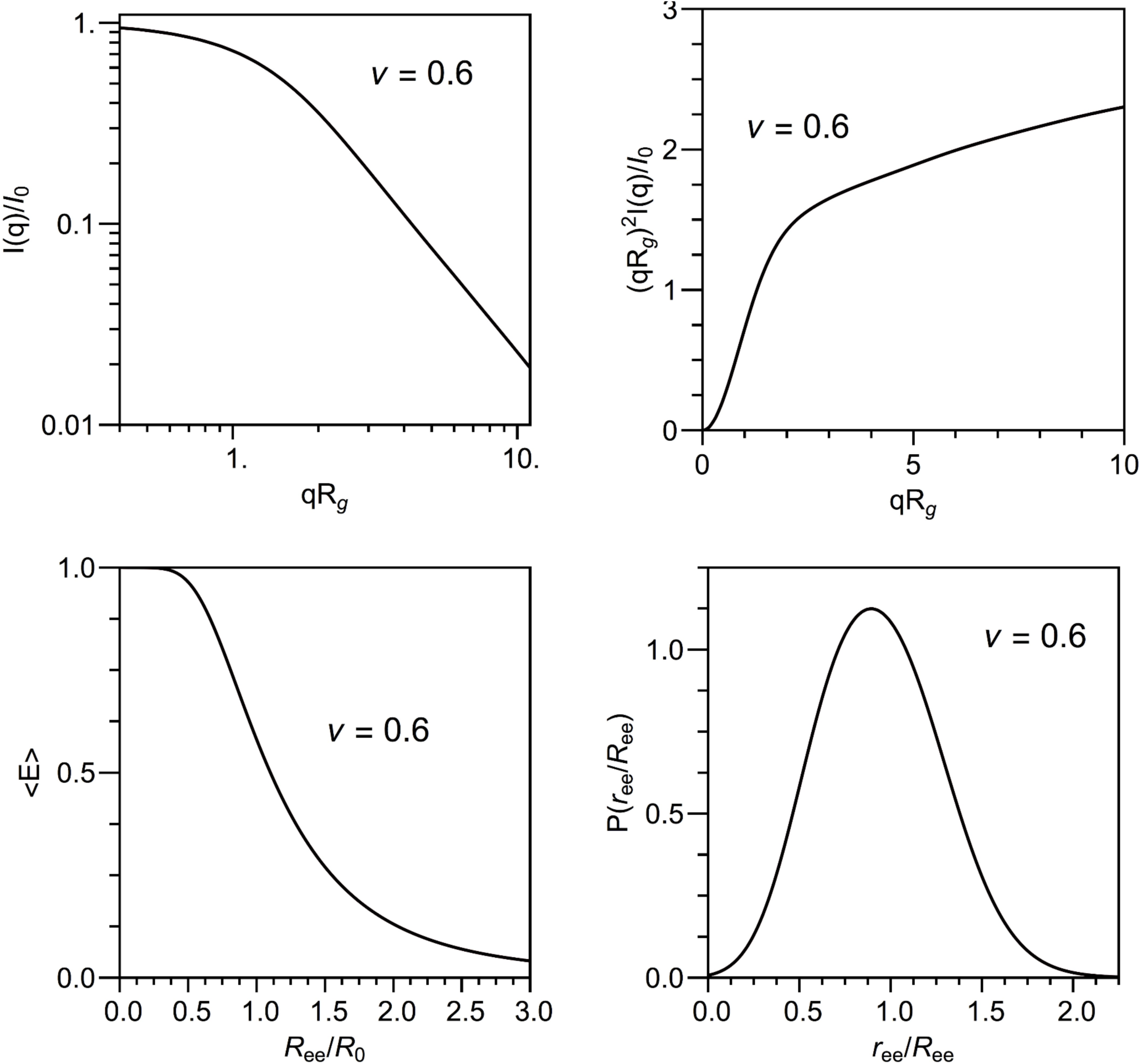
(Initial picture of movie found as attached file ‘MovieS1.avi’) Disordered homopolymer polymer functions for SAXS and FRET. Running video changes ν of black curve. Red curve is held at v=0.6. Note that the x-axis on the left plot is dimensionless and is thus scaled during fitting process.

**Movie S2:**
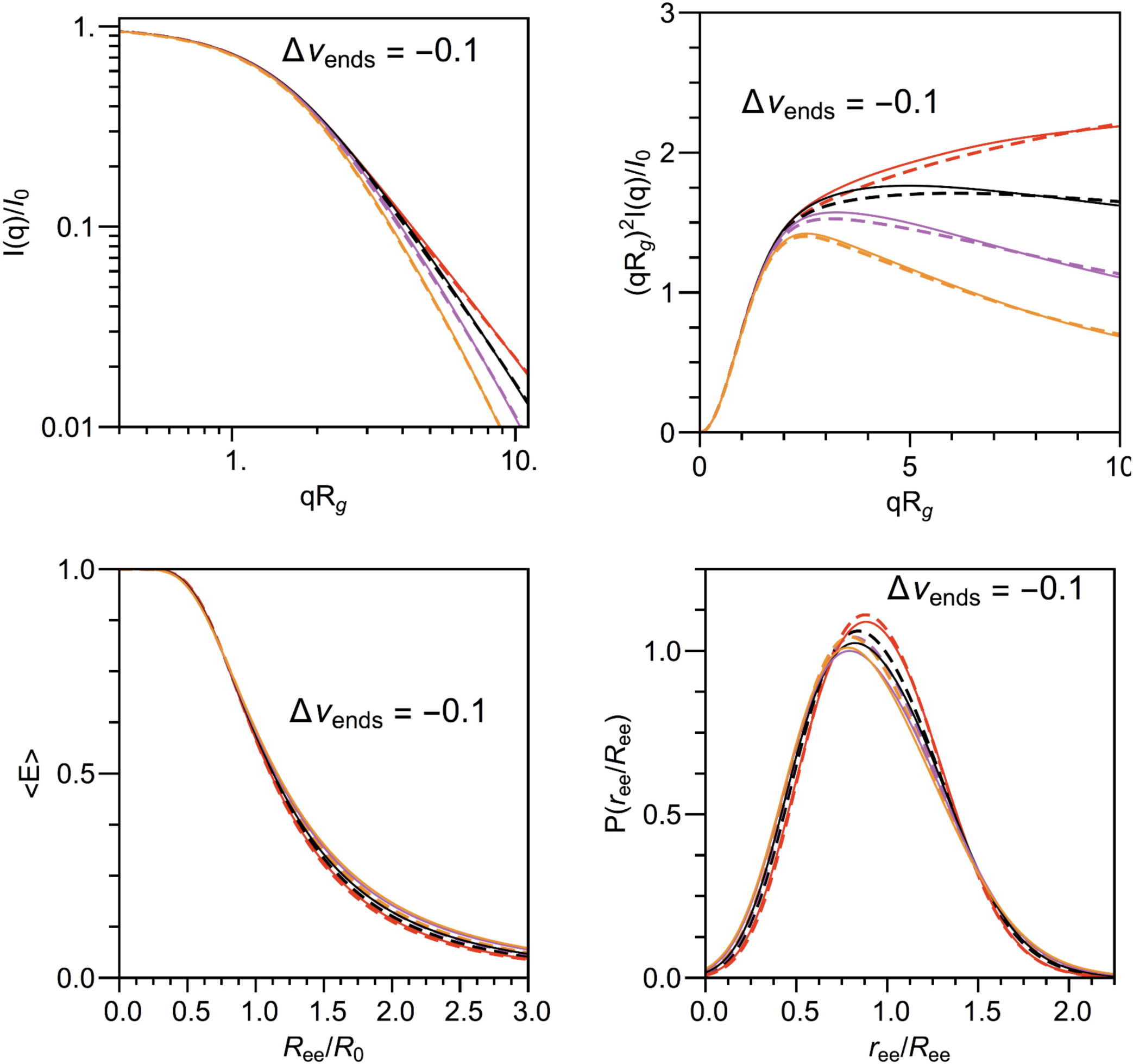
Disordered heteropolymer polymer functions for SAXS and FRET. (Initial picture of movie found as attached file ‘MovieS2.avi’) Running video changes Δv_ends_ of solid curve. Dashed curves are held at Δv_ends_=0.0. Red, black, purple, and orange curves are held at v=0.6, 0.55, 0.5, and 0.45, respectively. Note that the x-axis on the left plot is dimensionless and is thus scaled during fitting process.

## Footnotes

1 Error shown is the SEM of 3 measurements.

2 q is in units of inverse angstroms. When (59) is included in citation column, it provided data obtained from other studies.

